# Secretogranin-II plays a critical role in zebrafish neurovascular modeling

**DOI:** 10.1101/123125

**Authors:** Binbin Tao, Hongling Hu, Kimberly Mitchell, Ji Chen, Haibo Jia, Zuoyan Zhu, Vance L. Trudeau, Wei Hu

## Abstract

Neurons expressing *sgIIb* align with central arteries in hindbrain. We show that *sgIIb* is critical for neurovascular modeling the larval zebrafish mediated by MAPK and PI3K/AKT signaling *in vivo*.

**Abstract:** Secretoneurin (SN) is a neuropeptide derived from specific proteolytic processing of the precursor secretogranin II (SgII). In zebrafish and other teleosts there are 2 paralogs we previously named *sgIIa* and *sgIIb*. Our results showed that neurons expressing *sgIIb* were aligned with central arteries in hindbrain, demonstrating a close neurovascular association. Both *sgIIb*^-/-^ and *sgIIa*^-/-^ */sgIIb*^-/-^ mutant embryos were defective in hindbrain central artery development, while artery development in *sgIIa*^-/-^ mutant embryos was not affected. Hindbrain arterial and venous network identities were not affected in *sgIIb*^-/-^ mutant embryos, and the mRNA levels of Notch and VEGF pathway-related genes were not altered. However, the activation of MAPK and PI3K/AKT pathways were inhibited in *sgIIb^-/-^* mutant embryos. Injection of a synthetic SNb mRNA or delivery of the protein kinase activator N-arachidonoyl-L-serine could partially rescue the central artery developmental defects in the *sgIIb* mutants. This study provides the first *in vivo* evidence that *sgIIb* plays a critical role in neurovascular modeling the hindbrain.

## Introduction

The vertebrate vascular system is composed of highly branched networks of arteries, veins, and capillaries. The development of the vascular system occurs by two processes: vasculogenesis and angiogenesis. Vasculogenesis is the *de novo* assembly of the first blood vessels, whereas angiogenesis is the coordinated growth of endothelial cells (ECs) from the pre-existing vasculature (Risau, 1997). Both processes are essential for the maintenance of tissue growth and organ function in development (Carmeliet, 2003a). Numerous congenital or acquired diseases are associated with pathological vasculogenesis or angiogenesis (Carmeliet and Jain, 2000; Folkman, 1995; Psaltis and Simari, 2015). Among them, brain tumors, ischemic stroke and certain neurodegenerative diseases, including Alzheimer’s are associated with abnormal angiogenesis in the brain (Kim and Lee, 2009; Krupinski et al., 1994; Zlokovic, 2005). Therapeutic angiogenesis and anti-angiogenic therapies are regarded as promising strategies for treatment of such diseases (Chu and Wang, 2012; Jain, 2001; Wong et al., 2016). However, some of the larger clinical trials for therapeutic angiogenesis using single angiogenic factors have not corroborated the exciting early results (Krichavsky and Losordo, 2011). Combinations of multiple angiogenic factors to regulate angiogenesis may be a more promising approach. It is therefore necessary to uncover new pro-angiogenic and anti-angiogenic factors and to have a comprehensive understanding of the mechanism of angiogenesis.

Many ligands and their receptors are involved in the regulation of angiogenesis. These include Notch/Delta-like pathway (Siekmann and Lawson, 2007; Wang et al., 2014), VEGF pathway (Olsson et al., 2006), Wnt pathway (Dejana, 2010), shh pathway (Pola et al., 2001), angiopoietin (Suri et al., 1996), and netrins (Wilson et al., 2006) have been shown to be important. It is noteworthy that blood vessels are often aligned with nerves and display similar branching patterns. Some of these pathways or ligand-receptor complexes have been shown to act in parallel on both vascular and neural cells, demonstrating the interdependence and functional connection of the two systems sharing similar regulatory mechanisms (Larrivée et al., 2009). It may accelerate the discovery of mechanistic insights and new therapeutic opportunities to realize that vascular system and nervous system use some common genetic pathways (Carmeliet, 2003b).

Secretogranin-II (SgII), also known as chromogranin C, is mainly distributed in dense-core vesicles of many neurons and endocrine cells, and is overexpressed in some neuroendocrine tumors (Mahata et al., 1991). The neuropeptide secretoneurin (SN) is a short 31-43 conserved peptide derived from the larger ∼600 amino acid SgII precursor protein by prohormone convertase-mediated processing (Fischer-Colbrie et al., 1995). While several potential peptides may arise from SgII processing, SN is the only highly abundant neuropeptide with known biological activities (Fischer-Colbrie et al., 1995). Various physiological roles have been assigned to SN (Trudeau et al., 2012), including those related to reproduction (Zhao et al., 2009), neuroinflammation (You et al., 1996) and neurotransmitter release (Reinisch et al., 1993). In the rodent brain, SN is found predominantly in the phylogenetically older parts, overlapping partly but not completely with established neurotransmitter or peptidergic systems (Fischer-Colbrie et al., 1995; Trudeau et al., 2012). Supporting this, in the goldfish SN exhibits a more restricted CNS distribution mainly to the preoptic magnocellular neurons co-expressing nonapetides in the oxytocin/vasopressin family, certain hypothalamic nuclei, and posterior projections to hindbrain structures (Trudeau et al., 2012).

Until now, best described are the pro-angiogenic effects of SN. It has been reported that synthetic SN can promote capillary tube formation in human umbilical vein endothelial cells (HUVECs) *in vitro* (Kirchmair et al., 2004b). Synthetic SN can also induce neovascularization in the mouse cornea *in vivo*. The mechanism for the promotion of angiogenesis by SN may be not the same in different blood vessels. *In vitro* results show that the activation of mitogen-activated protein kinases (MAPK) by SN is dependent on vascular endothelial growth factor (VEGF) in human coronary artery endothelial cells (HCAECs) (Albrecht-Schgoer et al., 2012), but in HUVECs, the angiogenic effects by SN are VEGF-independent (Kirchmair et al., 2004b). To date, most of the studies on angiogenesis were carried out using exogenously applied SN. However, little is known about the effect of the endogenous SgII precursor protein or the SN neuropeptide on blood vessel formation as there are currently no *sgII* knockout animal models, and human mutations have not yet been identified. Moreover, the function of the *sgII* gene in the development of vertebrates is unknown. The aim of our study therefore was to investigate the role of *sgII* during early developmental stages.

In recent years, the zebrafish (*Danio rerio*) has emerged as an important vertebrate model system for studying angiogenesis in development (Chavez et al., 2016). The molecular mechanisms of zebrafish vessel formation are highly similar to those in humans and other vertebrates (Jinn et al., 2005). Zebrafish have the unique advantage for the study of brain angiogenesis because larvae can survive a few days without vascular development. External development, small brain, transparency of the zebrafish embryos and availability of endothelial specific fluorescent reporters offers unmatched optical access into the cerebral vasculature (Fujita et al., 2011; Lawson and Weinstein, 2002b).

In zebrafish and other teleost fishes, there exist two paralogous genes, *sgIIa* and *sgIIb* that generate SNa and SNb peptides, respectively (Zhao et al., 2010). In this study, we have generated the zebrafish *sgIIa*^-/-^, *sgIIb*^-/-^ and *sgIIa*^-/-^*/sgIIb*^-/-^ mutant lines using transcription activator-like effector nucleases (TALENs), and found *sgIIb*^-/-^ and *sgIIa^-/-^/sgIIb*^-/-^ mutants have specific defects in the development of hindbrain central arteries (CtAs). SgIIb plays a critical role in zebrafish neurovascular modeling that is mediated by MARK and PI3K/AKT signaling *in vivo*.

## Results

### *sgIIb* Is Expressed in The Central Nervous System of Zebrafish Embryos and *sgIIb*-Expressing Neurons and Central Arteries Are Aligned in Hindbrain

We have established the expression pattern of *sgIIb* in wild-type (WT) zebrafish embryos by RT-PCR and whole mount *in situ* hybridization (WISH) technique. *sgIIb* mRNA was first weakly detected by RT-PCR at 10 hours post-fertilization (hpf), its expression was increased over the 14-24 hpf period, then stabilized after 36 hpf (Fig. 1A). The WISH results reveal that *sgIIb* is mainly expressed in the central nervous system (CNS) at 24 and 36 hpf, and by 45 hpf *sgIIb* expression was concentrated in the brain (Fig. 1B). Then we examined the relationship between *sgIIb*-positive cells and *camk2d2* (calcium/calmodulin-dependent protein kinase (CaM kinase) II delta 2) expressing cells in hindbrain, Camk2d2 belongs to CaMKII family which functions in neuronal growth cone guidance and synaptic plasticity, among other functions (Mayford et al., 1995; Wen et al., 2004). Double-fluorescence *in situ* hybridization revealed the colocalization of *sgIIb* and *camk2d2* in hindbrain (Fig. 1C). To investigate the positional relationship between *sgIIb*-expressing cells and the vascular system in hindbrain, double-fluorescence *in situ* hybridization with *sgIIb* and vascular marker *kdrl* (kinase insert domain receptor-like) probes was performed in 36-45 hpf embryos. Our results show that the *sgIIb* expressing cells were aligned with the growing central arteries (CtAs) at 36-39 hpf (Fig. 1D-I, P, Q). At 42-45 hpf, *sgIIb* mRNA was widely distributed around CtAs in hindbrain (Fig. 1J-O, R). The expression of *sgIIa* was different from that of *sgIIb*. Firstly, *sgIIa* was detectable by 10 hpf (Fig. S1A), approximately 4 hr earlier than *sgIIb* (Fig. 1A). Thereafter, *sgIIa* expression increased gradually until it stabilized around 24 hpf. Expression of *sgIIa* was highly expressed in the forebrain, midbrain and ventral part of the neural tube (Fig. S1B). *SgIIa* was barely expressed in hindbrain (Fig. S1B), which contrasts significantly with the abundance of *sgIIb* transcripts in this region (Fig. 1B). Double-fluorescence WISH revealed that *sgIIa-po*sitive cells co-express the GABAergic neuron marker *tal2* (basic helix-loop-helix transcription factor) and *gad67*, the mRNA encoding the GABA-synthesizing enzyme glutamic acid decarboxylase 67 (Fig. S1C).

**Fig. 1.**
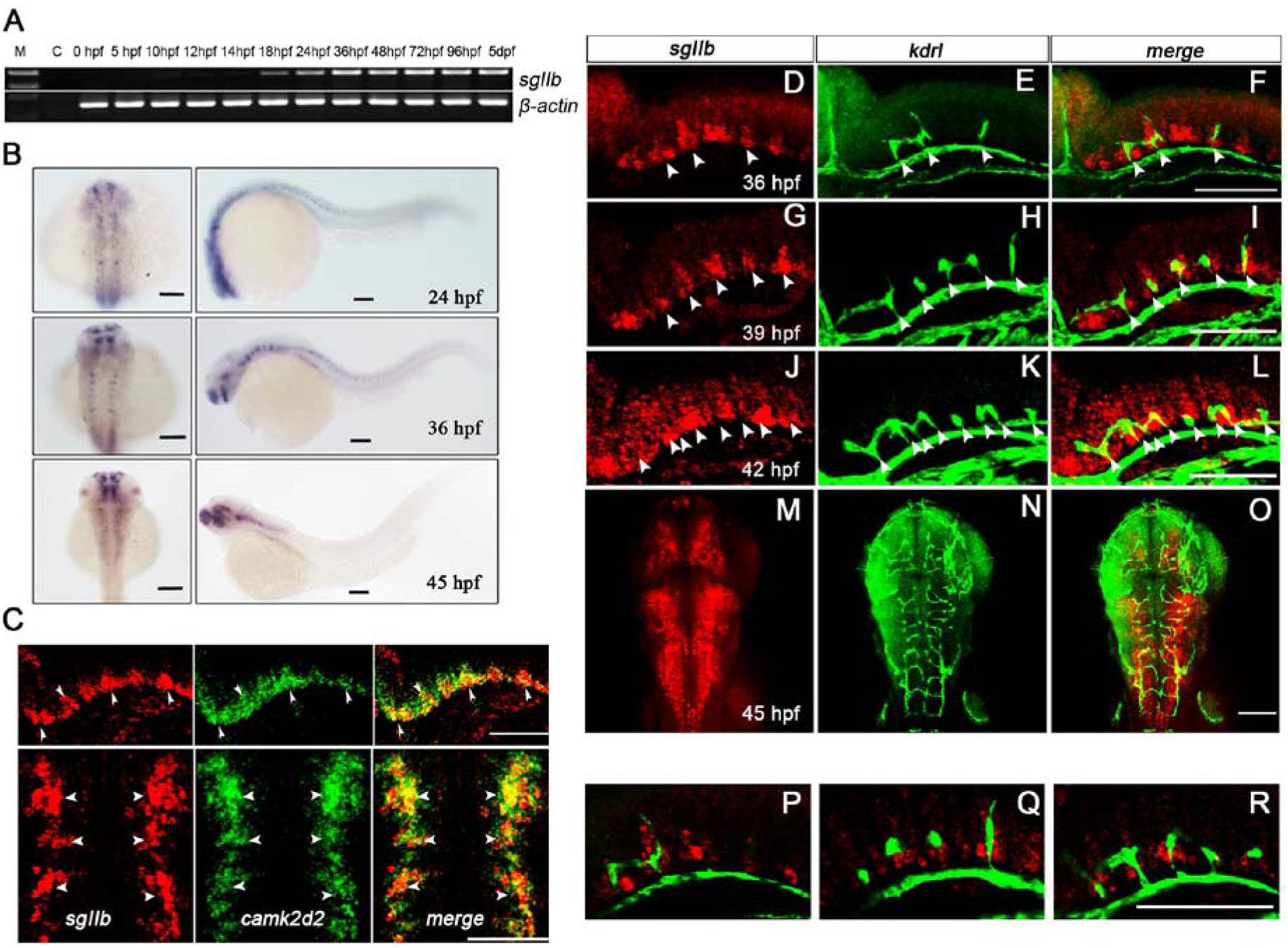
The Developmental Expression Pattern of *sgIIb* in The Central Nervous System of Zebrafish. **(A)** RT-PCR analysis for the temporal expression profile of *sgIIb* mRNA during embryogenesis and early larval developmental stages. M; molecular size marker, C; no template control, hpf; hours post-fertilization, dpf; days post-fertilization. **(B)** Whole-mount *in situ* hybridization of *sgIIb* in 24, 36 and 45hpf zebrafish embryos. **(C-R)** Mapping expression of *sgIIb* in relation to *camk2d2* and *kdrl* in the zebrafish hindbrain with double-fluorescence *in situ* hybridization. **(C)** Confocal imaging of *sgIIb* mRNA, *camk2d2* mRNA and the merged view at 36hpf. camk2d2-positive cells co-express *sgIIb* (arrowheads). **(D-O)** Maximal intensity projection of a confocal z-stack of *sgIIb* mRNA, *kdrl* mRNA and the merged view. kdrl-positive cells are aligned with *sgIIb* expressing cells (arrowheads). **(P-R)** Single confocal planes showing *sgIIb* and *kdrl* mRNA. Embryos were examined at 36 hpf **(D-F, P)**, 39 hpf **(G-I, Q)**, 42 hpf **(J-L, R)** and 45 hpf **(M-O)**. The scale bars represent 100 μm.

### Establishment of The Zebrafish *sgII* Mutant Lines with TALENs

To investigate the role of *sgII* during neurovascular development *in vivo*, two pairs of TALENs were designed for the zebrafish *sgIIa* and *sgIIb* genes. The *sgIIa* gene is located on chromosome 15 and *sgIIb* gene on chromosome 2, respectively. The TALEN target sites of *sgIIa* and *sgIIb* were both chosen following the ATG start site and in front of SNa and SNb domains (Fig. 2A, C). TALEN mRNAs were generated *in vitro* and injected into zebrafish embryos at the one-cell stage. The injected embryos were raised to adulthood (F0) and crossed with WT zebrafish, then the progeny (F1) were raised to adulthood and screened for *sgIIa* and *sgIIb* gene mutations. The *sgIIa* heterozygote with a 7-bp deletion and 2-bp insertion (-7,+2bp) or just a 7-bp deletion (-7bp) and the *SgIIb* heterozygote with a 7-bp insertion and a 5-bp (-5,+7bp) deletion or just a 10-bp deletion (-10bp) were screened out and further used to establish the *sgIIa*^-/-^ *and sgIIb*^-/-^homozygous mutant line (Fig. 2A, C). All of these mutant lines resulted in open reading frame-shift mutants of *sgIIa* or *sgIIb* gene, and thus generating truncated proteins with no functional SN peptides (Fig. 2B, D). The *sgIIb*^-/-^*/sgIIb*^-/-^ homozygous mutant line (*sgIIa*^-/-^-7,+2bp; *sgIIb*^-/-^-5,+7bp) was obtained by crossing the *sgIIa*^-/^*^-^* homozygote mutant line (-7,+2bp) with the *sgIIb*^-/-^homozygote (-5,+7bp). All crossings were performed using *in vitro* fertilization. Both qRT-PCR and WISH were used to detect mRNA levels of *sgIIa* and *sgIIb* in the mutant fish, and the data indicate that we have established stable zebrafish *sgII* mutant lines. The qRT-PCR results revealed that mRNA levels of *sgIIa* and *sgIIb* were also significantly decreased in *sgIIa* (Fig. 2E) and *sgIIb* (Fig. 2F) mutant embryos compared to WT, indicating a mechanism of nonsense-mediated mRNA decay (Chang et al., 2007). The WISH for both *sgIIa* and *sgIIb* (Fig. 2G) results confirmed the qRT-PCR data and indicate a loss of both *sgIIa* and *sgIIb* mRNAs. The pituitary is the major site of production of the SgIIa protein and SN peptide, so we monitored SN immunoreactivity in adults to determine the effects of mutations on protein production using a well-characterized polyclonal antibody (Zhao et al., 2006) that recognizes zebrafish SNa but not SNb (Fig. S2). Two bands corresponding to full-length precursor (∼62.2 KDa) and an intermediate fragment (∼59 KDa) were observed in WT zebrafish pituitaries. Our results reveal a lack of SgIIa precursor protein or any detectable proteolitically processed SNa-immunoreactive fragments in the adult pituitary gland of *sgIIa*^-/-^ mutant and *sgIIa*^-/-^*/sgIIb*^-/-^ double mutant fish. The ∼30 KDa SNa-immunoreactive fragment of SgIIa was dramatically increased in adult *sgIIb*^-/-^mutant fish (Fig. 2H), suggesting compensatory expression of *sgIIa* in adult *sgIIb*^-/-^mutant fish.

**Fig. 2.**
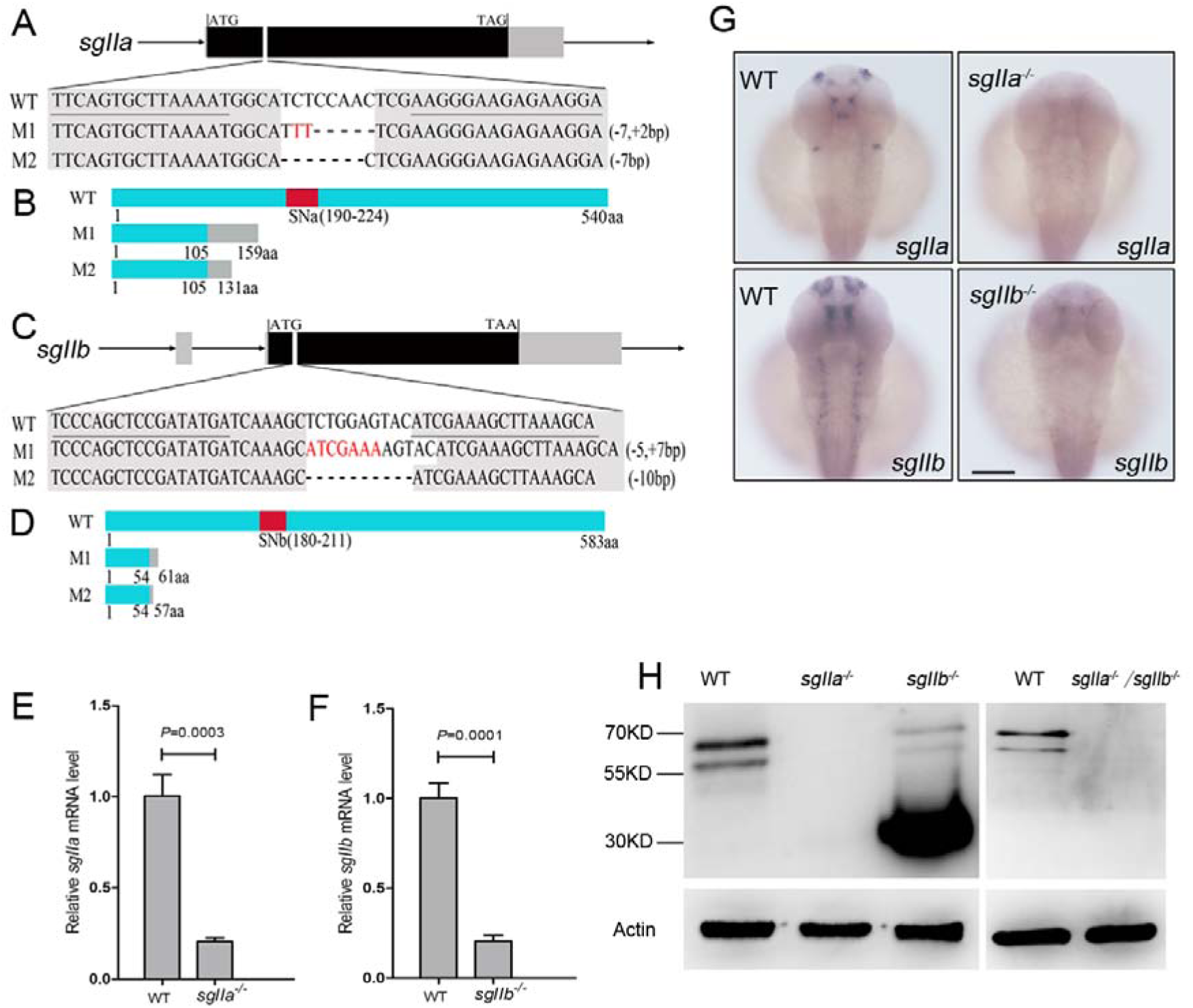
Targeted and Heritable Disruption of The *sgIIa* and *sgIIb* Genes in Zebrafish Using TALENs. **(A)** The location of the TALEN binding sites on the zebrafish *sgIIa* gene and two mutant lines of TALEN targeted *sgIIa* alleles. The TALEN binding sites are underlined. Deletions and insertions are indicated by dashes and red letters, respectively. **(B)** Schematic representation of the putative wild-type SgIIa protein and two mutated SgIIa proteins. **(C)** The location of the TALEN binding sites on the zebrafish *sgIIb* gene and two mutant lines of TALEN targeted *sgIIb* alleles. The TALEN binding sites are underlined. Deletions and insertions are indicated by dashes and red letters, respectively. **(D)** Schematic representation of the putative wild-type SgIIb protein and two mutated SgIIb proteins. (E and F) Relative mRNA level of *sgIIa* **(E)** or *sgIIb* **(F)** in *sgIIa*^-/-^ mutant or *sgIIb^-/-^* mutant 36 hours post-fertilization (hpf) embryos measured by qRT-PCR. Data shown are the mean ± SEM (3 independent experiments). Statistical significance was assessed using a two-tailed Student’s t-test. **(G)** Whole mount *in situ* hybridization of wild-type, *sgIIa*^-/-^mutant, *sgIIb*^-/-^mutant embryos. Antisense probes against *sgIIa* and *sgIIb* were visualized at 36 hpf. The scale bar represents 100 μm. **(H)** Western blotting analysis of the pituitary samples from wildtype, *sgIIa*^-/-^ mutant, *sgIIb*^-/-^ mutant and *sgIIa*^-/-^/*sgIIb*^-/-^ mutant adults at 120 days post-fertilization. A rabbit anti-goldfish SNa antiserum was used to detect the expression of SgII-related proteins containing the conserved central core of SN following published procedures (Zhao et al., 2006).

### Mutation of *sgIIb* Causes Defects of CtA Development In Vivo

The *sgIIa^-/-^, sgIIb^-/-^ and sgIIa^-/-^/sgIIb^-/-^mutant* zebrafish lines were crossed with the *Tg(kdrl:EGFP)* line that expresses green fluorescent protein (GFP) in the vascular system. Their progeny were raised to adulthood and intercrossed to establish 3 homozygous mutant lines with GFP expressed in vascular system: (*TG1*: *sgIIa^-/-^×Tg(kdrl:EGFP), TG2: sgIIb^-/-^×Tg(kdrl:EGFP), TG3:sgIIa^-/-^/sgIIb^-/-^×Tg(kdrl: EGFP)*). Central arteries are a set of vessels penetrating the hindbrain; they grow from the primordial hindbrain channels (PHBCs) to the basilar artery (BA) and interconnect the PHBCs and BA (Fig. 3A). *In vivo* confocal imaging results show that the specific defects of hindbrain CtA development in *TG2* and *TG3* embryos at 36, 39, 42, 45 and 48 hpf (Fig. 3B). Compared to the *Tg(kdrl:EGFP)* line, we found that the number of normal CtAs connected to the basilar artery (BA) was significantly decreased (Fig. 3C) while the number of disorganized CtAs increased (Fig. 3D) in *TG2* and *TG3* embryos at 45 and 48 hpf. The appearance of the CtA defects in *TG2* and *TG3* embryos were highly similar (Fig. 3C, D). In contrast, CtA development was not affected in *TG1* embryos (Fig. S3A). In addition, other blood vessels remained normal in *TG2* embryos (Fig. 4), demonstrating that *sgIIb* is critical for neurovascular modeling specifically in hindbrain. To examine the key roles of *sgIIb* in CtA development more precisely, we carried out time-lapse imaging from 36 to 45 hpf using *TG(kdrl:EGFP)* and *TG2*. Our results showed that most CtAs failed to sprout or connect to the BA in *TG2* embryos that lack SgIIb (Movie 1) compared with *TG(kdrl:EGFP)* (Movie 2). Since *sgIIa* is weakly expressed in hindbrain of zebrafish embryos (Fig. S1B), and the *sgIIa* mRNA level remained unchanged in *sgIIb* mutant embryos (Fig. S3B, C), *sgIIa* is unlikely to participate in the development of hindbrain CtAs.

**Fig. 3.**
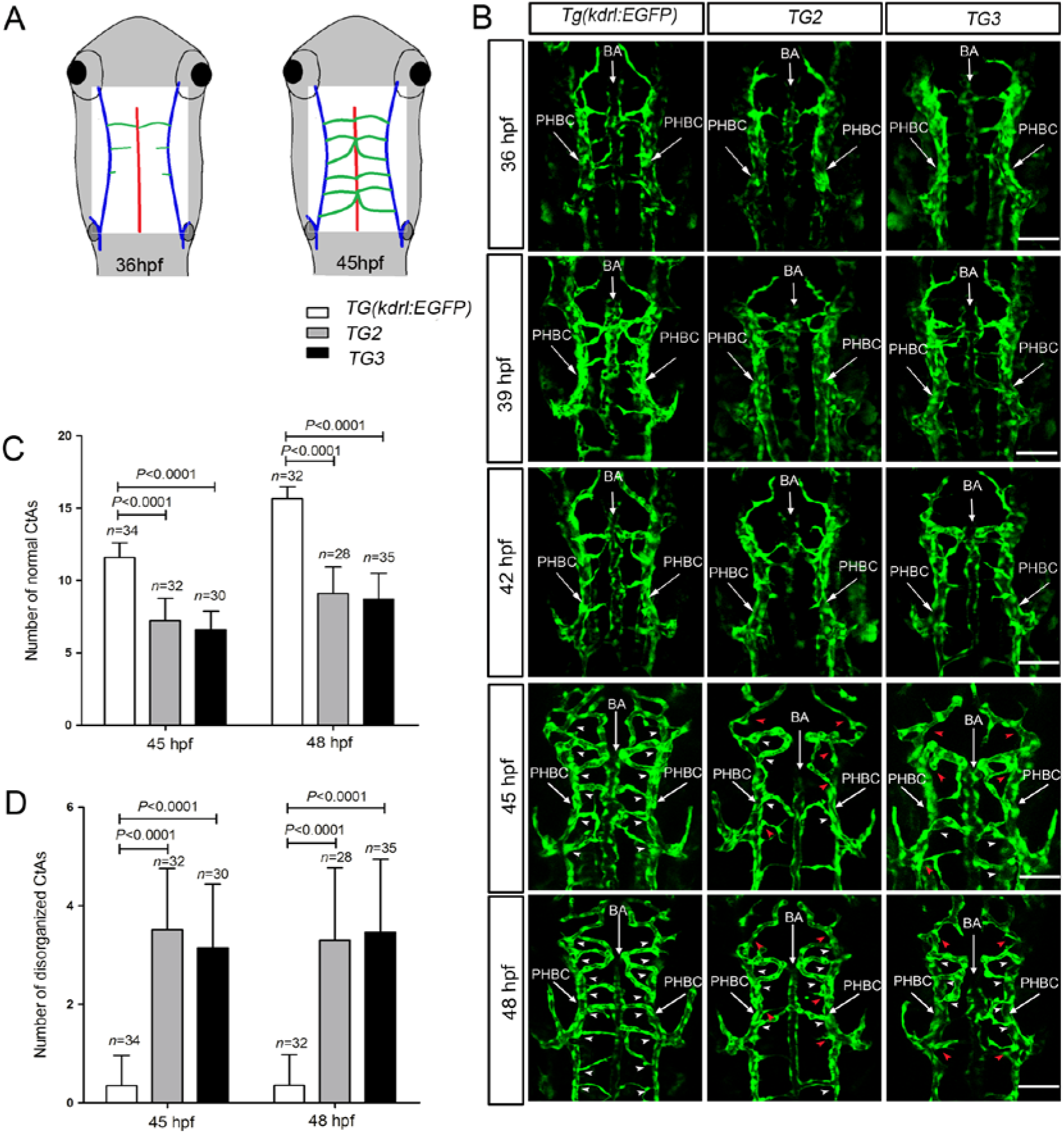
Mutation of *sgIIb* Causes Defects of CtA Development In Vivo. **(A)** Schematic vascular modeling in the hindbrain of 36hpf and 45 hpf wild-type zebrafish embryos. Primordial hindbrain channels (PHBC) are in blue, basilar artery (BA) is in red and the central arteries (CtA) are in green. **(B)** Maximal intensity projection of a confocal z-stack of *Tg*(*kdrl:EGFP*), *TG2: sgIIb*^-/-^×*Tg*(*kdrl:EGFP*) and *TG3: sgIIa*^-/-^*/sgIIb*^-/-^*×Tg*(*kdrl:EGFP*) zebrafish hindbrain. White arrowheads indicate normal CtAs connected to BA. Red arrowheads indicate disorganized CtAs. Embryos were examined at 36, 39, 42, 45 and 48 hours post-fertilization (hpf). The scale bars represent 50um. **(C and D)** Quantitative analysis of CtAs in *Tg*(*kdrl:EGFP*), *TG2* and *TG3* zebrafish hindbrains. Numbers of normal CtAs **(C)** and number of disorganized CtAs **(D)** were counted in *Tg*(*kdrl:EGFP*), *TG2* and *TG3* zebrafish hindbrain. Embryos were 45 hpf and 48 hpf. Data shown are the mean ± SEM (n = the number of embryos analysed). Statistical significance was assessed using a two-tailed Student’s t-test.

**Fig. 4.**
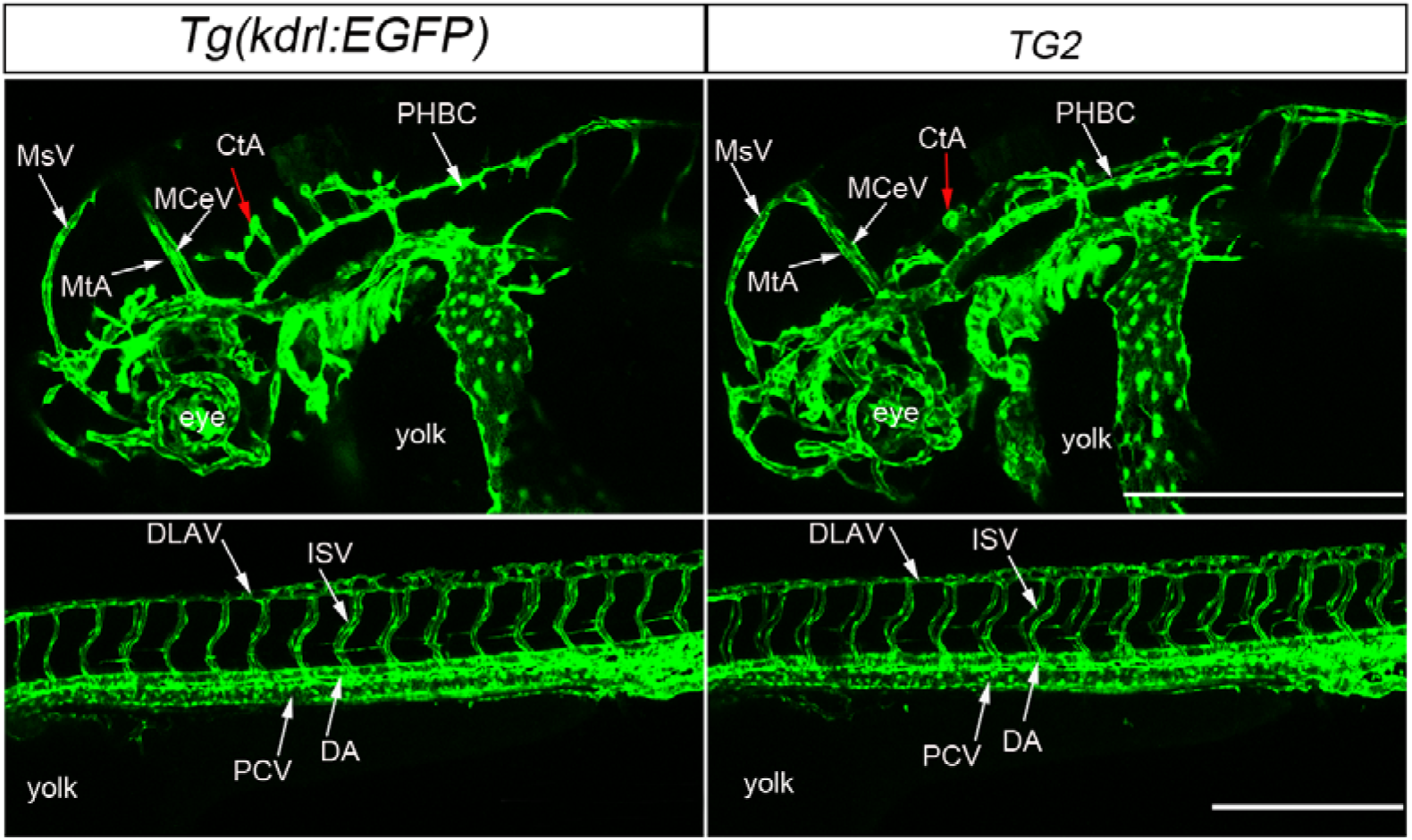
The Vasculature of *Tg*(*kdrl:EGFP*) and *TG2* Embryos at 45 Hours Post-Fertilization (Hfp). MsV: mesencephalic vein, MtA: metencephalic artery, MCeV: mid-cerebral vein, CtA: central artery, PHBC: primordial hindbrain channel, DLAV: dorsal longitudinal anastomotic vessel, ISV: intersegmental vessel, PCV: posterior cardinal vein, DA: dorsal aorta. Lateral views. The scale bars represent 25 μm.

Defects were also visualized using WISH for *cadherin 5* (*cdh5*), which is normally expressed in vascular endothelial cells. There was a clear reduction of *chd5* mRNA level in the hindbrain vasculature of 45 hpf *sgIIb*^-/-^ mutant embryos (Fig. 5A). This is consistent with the results observed in 45 hpf *TG2* or *TG3* embryos. A morpholino (MO) RNA targeting the start codon region of *sgIIb* (Fig. S4A) was synthesized and validated (Fig. S4B). Then *sgIIb*-MO knockdown experiments were performed in the *Tg*(*kdrl:EGFP*) line. At 45 hpf in the *Tg*(*kdrl:EGFP*) line *sgIIb* knockdown caused similar defects of CtA development (Fig. 5B) as noted with the *sgIIb* knockout experiments, while CtA development remained normal in *Tg*(*kdrl:EGFP*) embryos injected with a 5-mispair *sgIIb* MO. To date the only known bioactive peptide generated from the SgII precursor is SN, which is not generated in our mutants. We injected SNb mRNA into 1-cell embryos of the *TG2* line. Defects of CtAs development observed in *TG2* embryos could be partly rescued by SNb mRNA injection (Fig. 5C, D). Anti-phosphorylated histone H3 (PH3) antibody was used to detect the cells in M-phase of the cell cycle. Whole mount immunohistochemistry results show the number of proliferative tip ECs was decreased in the hindbrain CtAs of *sgIIb*^-/-^mutant embryos (Fig. 5E). Therefore, *sgIIb* is required for proliferation of tip ECs in zebrafish CtAs. To detect whether there are any neuronal phenotypes arising due to the deletion of *sgIIb*, double-fluorescence *in situ* hybridization were performed with *sgIIb* and *camk2d2* probes. Our results show that the *sgIIb* and *camk2d2* co-expressing neuronal cells were not affected in hindbrain of *sgIIb*^-/-^mutant embryos (Fig. 5F). These findings demonstrate the important role of *sgIIb* in neurovascular sprouting and pathfinding of zebrafish.

**Fig. 5.**
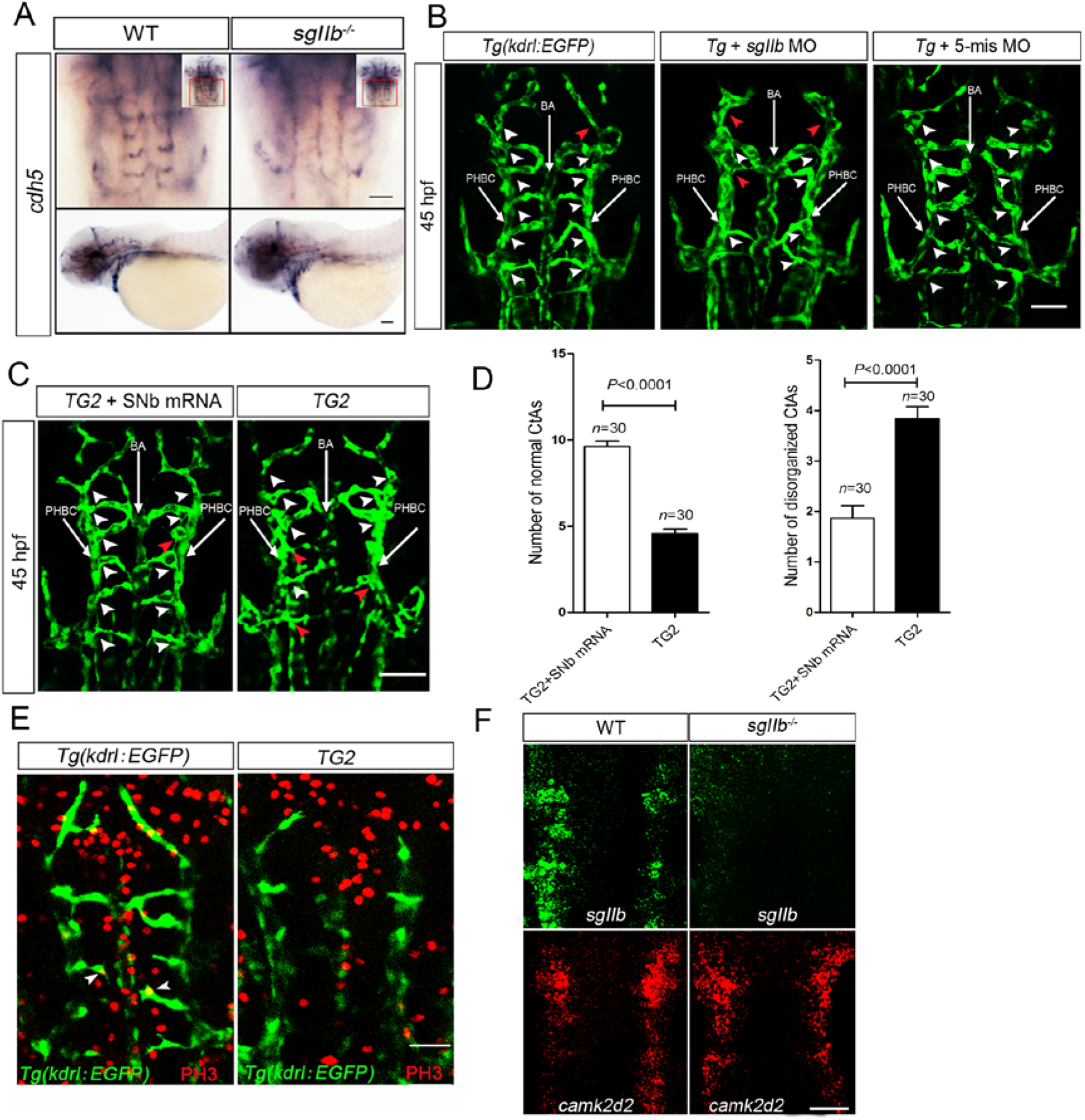
*sgIIb* Is Essential for CtA Development in Zebrafish Embryos. **(A)** Whole-mount *in situ* hybridization of the vascular marker *cdh5* in wild-type and *sgIIb^-/-^* mutant embryos. **(B)** Maximal intensity projection of a confocal z-stack of *TG2* embryos and *TG2* embryos injected with SNb mRNA. Red arrowheads indicate disorganized CtAs. **(C)** Maximal intensity projection of a confocal z-stack of *Tg*(*kdrl:EGFP*), *Tg*(*kdrl:EGFP*) injected with *sgIIb* MO and *Tg*(*kdrl:EGFP*) injected with sgIIb 5-mispair MO. White arrowheads indicate normal CtAs connected to BA. Red arrowheads indicate disorganized CtAs. BA: basilar artery. **(D)** Quantitative analysis of CtAs in hindbrains of *TG2* and TG2+SNb (*TG2* injected with SNb mRNA) zebrafish. Numbers of normal CtAs (left) and number of disorganized CtAs (right) were counted in 45 hpf *TG2* and *TG2*+ SNb zebrafish hindbraisn. Data shown are the mean ± SEM (n = the number of embryos analysed). Statistical significance was assessed using a two-tailed Student’s t-test. **(E)** Whole mount immunohistochemistry of 42hpf *Tg*(*kdrl:EGFP*) and *TG2* embryos with anti-phosphorylated histone H3 (PH3) antibody. **(F)** Double-fluorescence *in situ* hybridization of 36 hpf wild-type and *sgIIb*^-/-^ mutant embryos with *sgIIb* and *camk2d2* probes. The scale bars represent 50 μm.

### Mutation of *sgIIb* Does Not Affect Arterial-Venous Identity, and Notch and VEGF Pathways in Hindbrain

The establishment of arterial–venous identity is essential in the development of blood vessels (Lawson and Weinstein, 2002a), so we examined artery- and vein-specific markers in hindbrain of 36 hpf embryos. The expression of vein-specific markers (*dab2* and *flt4*) and artery-specific markers (*hey2, flt1* and *dll4*) were not affected in *sgIIb* mutant compared to WT embryos (Fig. 6A, B). It has been reported that Notch and VEGF pathways are critical for angiogenesis (Blanco and Gerhardt, 2013; Jakobsson et al., 2009). Fluorescence-activated cell sorting was used to obtain ECs from *Tg*(*kdrl:EGFP*) and *TG2* embryonic heads. The mRNA level of Notch and VEGF pathway-related genes was determined using qRT-PCR. As shown in Fig. 6C and Fig. S5, the mRNA levels of Notch pathway-related genes (including *notch1a, notch1b, notch2, dll4, hey2, hey1, hey6, dlc, dld*) and VEGF pathway-related genes (including *vegfaa, vegfab, vegfb, vegfc, vegfd, flt1, kdrl, flt4, nrpla, nrplb*) remained normal in ECs, indicating that Notch and VEGF pathways were not affected in the hindbrain of *sgIIb* mutant fish.

**Fig. 6.**
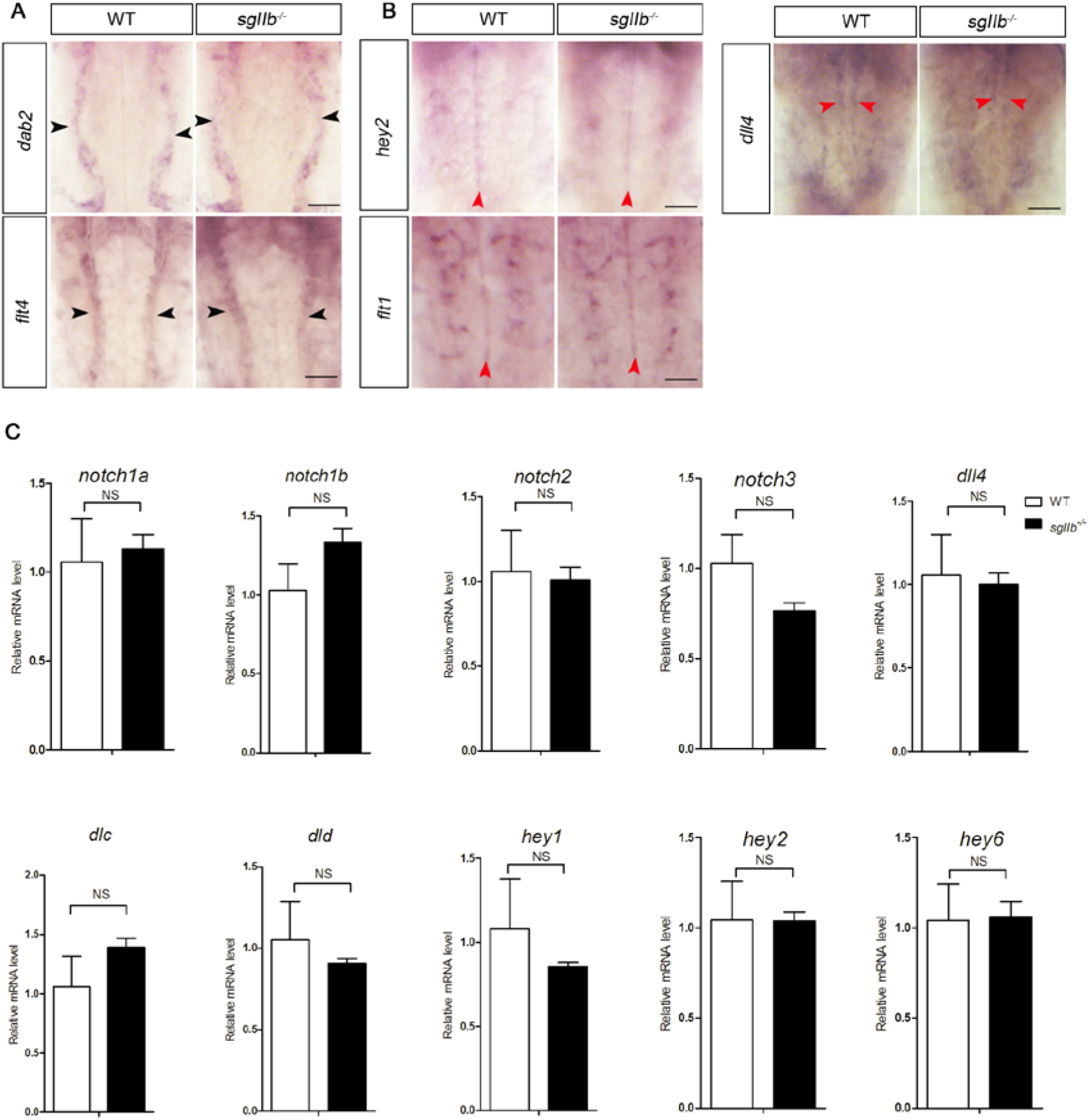
Mutation of *sgIIb* Does Not Affect Arterial-Venous Identity and The Expression of Notch Pathway-Related Genes. **(A and B)** Whole-mount *in situ* hybridization was performed in 36 hours post-fertilization (hpf) wild-type and *sgIIb* mutant zebrafish embryos. **(A)** *dab2* and *flt4* are venous markers and **(B)** *hey2, flt1* and *dll4* are arterial markers. Black arrowheads indicate the expression of venous markers, and red arrowheads indicate the expression of venous markers in hindbrain. The scale bars represent 50 μm. **(C)** The relative mRNA levels of *notch1b, notch1a, notch2, dll4, hey2, hey1, hey6, dlc, dld* were measured in FACS-sorted ECs from 36 hpf *Tg*(*kdrl:EGFP*) and *TG2* embryonic heads using qRT-PCR. Data shown are the mean ± SEM (3 independent experiments). Statistical significance was assessed using a two-tailed Student’s t-test. NS, not significant.

### The Role of *sgIIb* in Neurovascular Modeling Is Mediated by MAPK and PI3K/AKT Signaling In Vivo

Previous studies have demonstrated that MAPK and PI3K/AKT pathways participate in the proliferation and migration of ECs (Karar and Maity, 2011; Mavria et al., 2006). We examined the MAPK and PI3K/AKT pathways in *sgIIb* mutant embryos with defects of CtA development. The expression levels of p-ERK1/2 and p-AKT in *sgIIb* mutant embryos were much lower than in WT, while the levels of total-ERK1/2 and total-AKT were unchanged (Fig. 7A, B). These results suggested that the activation of MAPK and PI3K/AKT were inhibited in the *sgIIb* mutants. Accordingly, when we injected *SNb* mRNA into 1-cell *sgIIb* mutant embryos, the expression levels of p-EK1/2 and p-AKT were increased at 36 hpf (Fig. 7C), while the levels of total-ERK1/2 and total-AKT remained unchanged. N-arachidonoyl-L-serine is a known MAPK and PI3K/AKT pathways activator (Milman et al., 2006). Delivery of ARA (50 μM) increased p-EK1/2 and p-AKT level in *sgIIb* mutant embryos (Fig. 7D). Furthermore, CtA developmental defects in *TG2* fish could be partially rescued by ARA treatment (Fig. 7E, F).

**Fig. 7.**
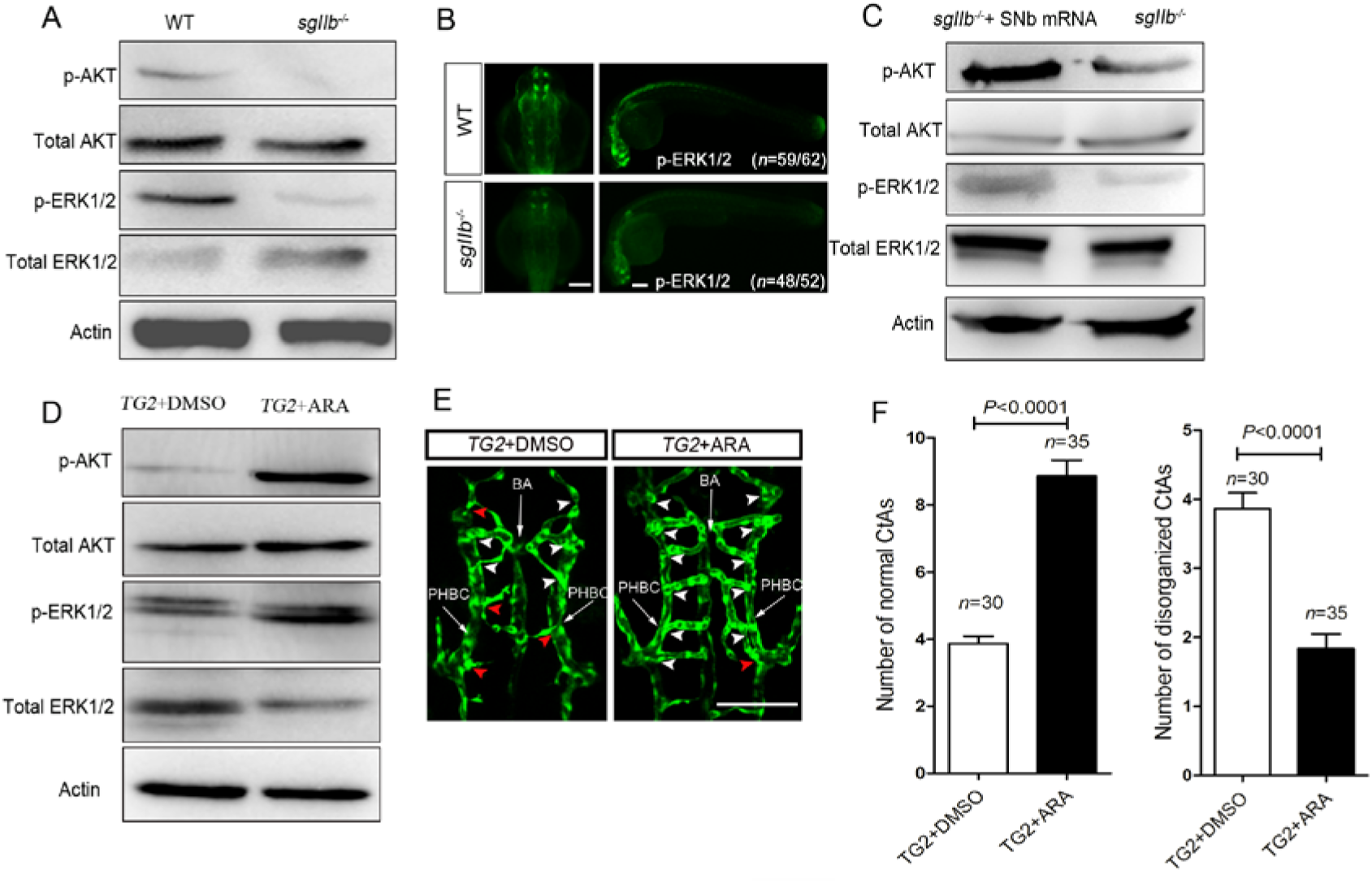
Mutation of *sgIIb* Inhibited The Activation of MAPK and PI3K/AKT In Vivo. **(A)** Western blots of phospho-AKT (p-AKT), total AKT, phospho-ERK1/2 (p-ERK1/2), total ERK1/2 expression in 36 hpf wild-type and *sgIIb*^-/-^ mutant embryos. Actin was used as a loading control. **(B)** Whole-mount immunohistochemistry of phospho-ERK1/2 (p-ERK1/2) expression in 36 hours post-fertilization (hpf) wild-type and *sgIIb*^-/-^ mutant embryos. The frequency of embryos with the indicated phenotypes is shown in the bracket of each group. **(C)** Western blots of phospho-AKT (p-AKT), total AKT, phospho-ERK1/2 (p-ERK1/2), total ERK1/2 expression in 36 hpf *sgIIb*^-/-^ mutant and *sgIIb*^-/-^ mutant embryos injected with *SNb* mRNA. Actin was used as a loading control. **(D)** Western blots of phospho-AKT (p-AKT), total AKT, phospho-ERK1/2(p-ERK1/2), total ERK1/2 expression in 36 hpf *sgIIb*^-/-^ mutant embryos incubated with DMSO (control) or N-arachidonoyl-L-serine (ARA; 50 μM). Actin was used as a loading control. **(E)** Maximal intensity projection of a confocal z-stack of 45 hpf *TG2* (*sgIIb-/-×Tg*(*kdrl:EGFP*) embryos incubated with DMSO or ARA. White arrowheads indicate normal CtAs connected to BA. Red arrowheads indicate disorganized CtAs. **(F)** Quantitative analysis of CtAs in *TG2* + DMSO (*TG2* exposed to DMSO) and *TG2* + ARA (*TG2* exposed to ARA) zebrafish hindbrains. Numbers of normal CtAs (left) and number of disorganized CtAs (right) were counted in 45 hpf *TG2* + DMSO and *TG2* + ARA zebrafish hindbrains. Data shown are the mean ± SEM (n = the number of embryos analysed). Statistical significance was assessed using a two-tailed Student’s t-test. The scale bars represent 100 μm.

## Discussion

In this study, we have provided the first *in vivo* evidence that *sgIIb* plays a critical role in neurovascular modeling of the hindbrain that is mediated by activation of the MAPK and PI3K/AKT pathways. The neuropeptide SN, processed from SgII, has multiple physiological functions (Albrecht-Schgoer et al., 2012; Trudeau et al., 2012; Zhao et al., 2009). It has been reported that synthetic SN can promote angiogenesis *in vitro* and *in vivo* (Kirchmair et al., 2004b). Therefore, the function of SN in angiogenesis has attracted considerable interest in recent years, yet the role of SN in CNS-specific angiogenesis has remained unknown until now. To our knowledge, no other early developmental roles for *sgII* have been reported previously.

To investigate the role of *sgII* in CNS-specific angiogenesis, we generated *sgII* mutant zebrafish lines with GFP expressed in the vascular system. Our *in vivo* confocal imaging results showed that the number of normal CtAs was decreased while the number of disorganized CtAs was increased in the hindbrain of *sgIIb* mutant embryos. Time-lapse imaging indicated that depletion of *sgIIb* alone in zebrafish could lead to severe defects of CtA sprouting and pathfinding. While neurons co-expressing *sgIIb* and *camk2d2* were not affected in *sgIIb* mutant embryos. Injection of SNb mRNA could partly rescue the defects of CtA caused by the *sgIIb* mutation. It is thus demonstrated that *sgIIb* participates in neurovascular modeling. Despite the different physiological functions of the vascular and nervous systems, blood vessels are often aligned with nerves, and the two systems share some common molecular mechanisms (Carmeliet and Tessier-Lavigne, 2005; Park et al., 2004). Axon guidance molecules including netrins, ephrins, slits, semaphorins and their receptors are cues in vascular morphogenesis (Carmeliet and Tessier-Lavigne, 2005). They participate in blood vessel navigation by either repelling or attracting the filopodia of tip endothelial cells. We found that *sgIIb* is mainly expressed in CNS in zebrafish embryos and co-expressed with *camk2d2*, one member of CaMKII family which functions in growth cone guidance. The sgIIb-expressing cells were aligned with CtAs in hindbrain. Defects of CtA sprouting and pathfinding in *sgIIb* mutant embryos imply that *sgIIb* is involved in vascular guidance in hindbrain. These data suggest that SgIIb may act as a guidance cue in neurovascular modeling. Our results demonstrate that *sgIIb* plays an important role in neurovascular modeling in zebrafish.

Notch and VEGF signaling are fundamental for angiogenesis in physiological and pathological conditions. It has been demonstrated that VEGF signaling is stimulatory for angiogenic sprouting, while Notch signaling is inhibitory (Covassin et al., 2006b; Suchting et al., 2007; Zhong et al., 2001; Zhong et al., 2000). In hindbrain, Notch and VEGF signaling also play important roles in arterial-venous identity, which is regarded as an essential process of vascular development (Bussmann et al., 2011; Lawson and Weinstein, 2002a). We found that the expression of Notch and VEGF pathway-related genes were not affected in *sgIIb* mutant brain ECs and arterial-venous identity of hindbrain was normal. Immunohistochemistry with an anti-PH3 antibody indicated that the proliferation of tip ECs was decreased in *sgIIb* mutant embryos. As we know, MAPK system and PI3K/AKT pathway play important roles in the proliferation and migration of ECs. Using loss-of-function and gain-of-function methods, we found that MAPK and PI3K/AKT pathways were inhibited in *sgIIb* mutant embryos *in vivo*. Injection of SNb mRNA or delivery of the MAPK and PI3K/AKT pathways activator ARA could restore their function and partially rescue CtA developmental defects in *sgIIb* mutant embryos. This is consistent with previous *in vitro* reports showing that culturing HUVECs and endothelial progenitor cells (EPCs) with SN could activate the MAPK system and the PI3K/AKT pathway (Kirchmair et al., 2004a; Kirchmair et al., 2004b). Our results indicate that the MAPK system and PI3K/AKT pathway are mediators of *sgIIb* effects on neurovascular development. However, there are conflicting reports whether the activation of the MAPK system and PI3K/AKT are dependent on VEGF. In HUVECs, the activation of MAPK and PI3K/AKT pathway by SN is not dependent on VEGF (Kirchmair et al., 2004b). Results obtained from HCAECs indicate that SN could promote binding of VEGF to its co-receptors heparin and neuropilin-1, demonstrating the role of SN in VEGF signaling (Albrecht-Schgoer et al., 2012). Our data provide evidence that VEGF pathway-related genes and VEGF-dependent arterial-venous identity were not altered in *sgIIb*^-/-^ mutants.

In conclusion, we provide evidence that *sgIIb* plays an important role in neurovascular modeling in zebrafish. Cells expressing *sgIIb* were found in close association with vascular markers in the hindbrain, so this is the likely source of SgIIb and/or SNb that controls the development of these CtAs. Ischemic stroke and neurodegenerative diseases, for example, Alzheimer’s disease, are associated with abnormal angiogenesis in the brain. It is significant that SN promotes neuroprotection in mouse models of stroke (Shyu et al., 2008) and that proteomic markers for SgII are significantly decreased in CSF in Alzheimer patients (Spellman et al., 2015). Here, we show that injection of SNb mRNA could rescue the CtA defects in *sgIIb* mutant embryos. Targeting the SgII system may therefore represent a new avenue for the treatment of vascular defects in the CNS.

## Materials and Methods

### Zebrafish Care and Maintenance

AB strain zebrafish (*Danio rerio*) and their embryos were raised and maintained under standard conditions at 28.5°C. Embryos were staged as previously described (Kimmel et al., 1995). *Tg*(*kdrl:EGFP*) zebrafish were obtained from the China Zebrafish Resource Center (CZRC, Wuhan, China). The experiments involving zebrafish were performed under the approval of the Institutional Animal Care and Use Committee of the Institute of Hydrobiology, Chinese Academy of Sciences.

### Establishment of Zebrafish Mutant Lines

The paired TALENs for *sgIIa* or *sgIIb* were constructed using the golden gate method as described previously (Cermak et al., 2011; Liu et al., 2014). The final TALEN plasmids were linearized using Not1. TALEN mRNAs were synthesized using the mMessage mMACHINE SP6 Kit (Ambion, Inc., Austin, TX) and purified by LiCl precipitation. To generate zebrafish mutant lines, 250-500 pg TALEN mRNAs were microinjected into one-cell stage wild-type zebrafish embryos. Injected embryos were raised to adulthood and then outcrossed with WT fish to identify founders that transmitted mutations through the germ line. Mutations were genotyped by competitive PCR (Liu et al., 2014) and confirmed by sequencing.

### RNA Synthesis and Knockdown Experiments

For making SNb mRNA, the zebrafish *sgIIb* fragment encoding SNb was cloned into RNA synthesis plasmid pCS2+, and Capped mRNA was synthesized with the mMessage mMachineSP6 Kit (Ambion, Inc., Austin, TX) and purified by LiCl precipitation. The *sgIIb* specific antisense morpholino oligonucleotides (*sgIIb* MO5’-GTTTGGGTAGCGACAACATCATACC-3’) and the *sgIIb* 5-base mismatch morpholino oligonucleotides (5-mis *sgIIb* MO 5’-GTTaGcGTAGCcACAAgATgATACC-3’) were designed and synthesized by Gene Tools (Philomath, OR, USA), dissolved to 1mM in nuclease-free water and stored at -20°C. For microinjection, MOs were diluted in 1×Danieau’s buffer (58 mM NaCl, 0.7 mM KCl, 0.4 mM MgSO4, 0.6 mM Ca(NO3)2, 5 mM HEPES, pH 7.6) supplemented with phenol red. The effectiveness of the *sgIIb* MO was tested by co-injecting MO with the pEGFP-N1 plasmid fused in frame with the MO target site. For some experiments, the morphant and control embryos at 24 hpf were incubated in fresh egg water supplemented with 50μm ARA or 0.5% DMSO.

### Embryo Mounting, Confocal Microscopy and Image Processing

For *in vivo* confocal imaging, embryos were mounted in 1 % low-melt agarose after anaesthetized in 168 mg/l tricaine. Fluorescence photomicrographs were collected with laser scanning confocal microscope (Zeiss LSM710). Time-lapse movies were prepared using the software program ZEN (Zeiss). Egg water containing tricaine and propylthiouracil was warmed to approximately 28 °C and circulated during the observation period, Z stacks were collected at 10 min intervals. Images were processed using Adobe Photoshop CS3 Extended. The number of normal CtAs and the number of disorganized CtAs were counted using a fluoroscope (Leica M250). The unpaired two-tailed Student’s t-test as implemented in GraphPad Prism 5 was used to analyze the data.

### Whole Mount In Situ Hybridization and Double Fluorescence In Situ Hybridization

*sgIIa, sgIIb, camk2d2, kdrl, cdh5, dab2, flt1, flt4, hey2, dll4, tal2, gad67* sequences were cloned by PCR amplification, PCR products were subcloned into the pMD18-T vector (TAKARA). Digoxigenin labeled antisense probes for *sgIIa, sgIIb, cdh5, dab2, flt1, flt4, hey2, dll4* were synthesized using DIG RNA Labeling Mix (Roche Diagnostics, Mannheim, Germany). Whole mount in situ hybridization was carried out as described previously (Thisse and Thisse, 2008). Fluorescein-labeled antisense probes for *camk2d2, kdrl, tal2, gad67* were synthesized using fluorescein RNA Labeling Mix (Roche Diagnostics, Mannheim, Germany). Double fluorescence in situ hybridization was performed as described (Lauter et al., 2011). Primers used in whole mount *in situ* hybridization can be found in Table S1.

### Semi-Quantitative Reverse-Transcriptase PCR (RT-PCR) and Real-Time Quantitative PCR (RT-qPCR)

Heads of 36hpf zebrafish embryos were cut off using a pair of forceps. Endothelial cell (ECs) dissociated from 300-400 heads of *Tg*(*kdrl: EGFP*) and *TG2*(*sgIIb^-/-^×Tg*(*kdrl:EGFP*)) were sorted using the FACSAria™ III (BD Biosciences) as described previously (Covassin et al., 2006a). Total RNA was extracted from the sorted ECs or 40 zebrafish embryos (WT, *sgIIa*^-/-^, *sgIIb*^-/-^, *sgIIa*^-/-^*/sgIIb*^-/-^) at desired stages using TRIzol reagent (Invitrogen) and resuspended in diethyl-pyrocarbonate (DEPC)-treated water. The quality of extracted RNA was confirmed by UV spectrophotometer and agarose gel electrophoresis. Total RNA was reverse-transcribed into cDNA using ReverTra Ace M-MLV (TOYOBO) with random primers. For RT-PCR, the cDNA samples were PCR-amplified using gene-specific primers as listed in Table S1 (supplementary material). RT-qPCR was carried out on a Roche LightCycler 480 real-time PCR system using 2×SYBR green real-time PCR mix (TOYOBO), all values were normalized to the level of β-actin mRNA, primers used in RT-PCR (Table S2) and RT-qPCR (Table S3) can be found in supplementary material.

### Western Blotting

Embryos were de-yolked as described (Link et al., 2006). The Total Protein Extraction Kit (Sangon Biotech, China) was used to lyse zebrafish embryos and adult zebrafish pituitaries according to the manufacturer’s instructions, the proteins of each lysate were separated by 12.5% sodium dodecyl sulfate polyacrylamide gel electrophoresis (SDS-PAGE) and trans-blotted onto nitrocellulose membrane (Millipore), probed with indicated primary antibodies against phospho p44/42 MAPK (phospho-Erk1/2, 1:2000), p44/42 MAPK (total Erk1/2, 1:1000), p-AKT(1:2000), total AKT (1:1000) (all from Cell Signaling Technology), SNa (1:1000) (Zhao et al., 2006), β-actin (Bioss Company, China, 1:1500 diluted), then the membrane was washed in phosphate buffered saline containing Tween-20 (PBST) for 4×5 min, incubated with horseradish peroxidase-conjugated goat anti-rabbit or goat anti mouse secondary antibodies and detected using SuperSignal West Pico Chemiluminescent Substrate (Thermo Scientific). The signals were visualized using ImageQuant LAS 4000 mini system (GE Healthcare).

### Whole Mount Immunohistochemistry

Whole mount immunohistochemistry was performed using phosphor p44/42 MAPK (phospho-Erk1/2, 1:1000, CST) as primary antibody and DyLight 488 goat anti-mouse IgG (Abbkine, USA, 1:200) as secondary antibody. Briefly, dechorionated zebrafish embryos were fixed in 4% PFA in phosphate-buffered saline (PBS) overnight at 4°C, then embryos were dehydrated in 100% methanol for 15 min at room temperature and stored at -20°C in 100 % methanol for at least two hours before use. Embryos were permeabilized in acetone (-20°C) for 10 minutes, washed with phosphate-buffered saline containing TritronX100 (PBST), and incubated with the primary antibody overnight at 4°C, then washed extensively and incubated with the secondary antibodies overnight at 4°C. Embryos were washed extensively again and mounted in glycerol

### Statistical Analysis

Statistical analysis was performed using unpaired two-tailed Student’s t-test as implemented in GraphPad Prism 5 (GraphPad Software Inc., San Diego, CA, USA). Data are presented as means ± SEM.

## Author contributions

W.H., V.L.T., Z.Z., and B.T. designed the research; B.T., H.H., and K.M. developed methods and performed research; W.H., Z.Z., and J.C. contributed new reagents/analytic tools; B.T., W.H., H.J., and V.L.T. analyzed data; and B.T., V.L.T., and W.H. wrote the manuscript.

The authors declare no conflict of interest.

## Acknowledgements

This work was supported by the National Natural Science Foundation of China (Grant No. 31325026 to WH). Funding from the Natural Sciences and Engineering Research Council of Canada (to VLT) and the University of Ottawa International Research Acceleration Program (to VLT and WH) is acknowledged with appreciation. We are grateful to Ms. Fang Zhou for providing confocal services (Analytical & Testing Center, IHB, CAS), and Professors Marie-Andrée Akimenko (University of Ottawa) and Jingwei Xiong (Peking University) for their valuable discussion and critical reading of the manuscript.

**Secretogranin-II plays a critical role in zebrafish neurovascular modelling**

Binbin Tao, Hongling Hu, Kimberly Mitchell, Ji Chen, Haibo Jia, Zuoyan Zhu, Vance L. Trudeau, and Wei Hu

